# Extracellular matrix proteoglycans lumican and biglycan promote innate immune signals and viral clearance in HSV-1 infections of mouse corneas

**DOI:** 10.64898/2026.09.15.751741

**Authors:** George Maiti, Madhuri Koduri, Elizabeth Shin, Yoonchul Shin, Trisha Sinha, Alireza Khodadadi-Jamayran, Hongmin Yun, Anthony J. St. Leger, Shukti Chakravarti

## Abstract

The extracellular matrix (ECM) regulates innate immunity, but its role in HSV-1 keratitis is unclear. Here, we identify the ECM proteoglycans lumican (Lum) and biglycan (Bgn) as key regulators of early antiviral defense in the cornea. *Lum^-/-^* and *Bgn^KO^*mice showed impaired viral clearance, increased corneal opacity, and worse keratitis than wild-type controls. These defects were associated with reduced early recruitment of neutrophils, monocytes, and plasmacytoid dendritic cells, and blunted induction of pro-inflammatory cytokines, chemokines, and type I interferons. In infected wild-type corneas, Lum and Bgn were upregulated, including in the basal epithelium, where Ccl2 emerged as an early epithelial response that was diminished in knockout mice. In human corneal epithelial cells, recombinant LUM or BGN enhanced HSV-1-induced CCL2 secretion, and human corneal organoids recapitulated epithelial viral tropism and CCL2 induction. Together, these findings identify Lum and Bgn as regulators of epithelial-innate immune crosstalk that promote early antiviral immunity during corneal HSV-1 infection.

## Introduction

Herpes simplex virus-1 (HSV-1) is a double-stranded DNA virus that can cause epithelial or stromal keratitis or both ^1^. While clinical symptoms from primary infections in adult humans are rare, reactivation of the virus in the trigeminal ganglion and re-entry in the cornea cause inflammatory epithelial and stromal keratitis. Vision loss is driven not only by viral cytopathic effects but also by host inflammatory responses that disrupt corneal transparency. The immune-restricted cornea engages robust early innate immune signals in epithelial cells, stromal keratocytes, and resident immune cells to control inflammation and preserve optical transparency ^2^. Primary infection of mouse corneas is often used to investigate host responses to viral presence in the cornea. The corneal epithelial and stromal cells detect viral nucleic acids and structural components through pattern recognition receptors, including Toll-like receptors (TLRs), and trigger signaling networks that induce pro-inflammatory cytokines like IL-1β, IL-6, TNF, anti-viral type I interferons (IFNs), and chemokines IL-8 (humans), CXCL1 (mice), CCL2, CCL5, and CXCL10^3–8^. These cytokines and chemokines signal the recruitment of neutrophils, inflammatory monocytes, macrophages, and dendritic cells (DCs), which together contribute to viral control and subsequent adaptive immune priming^8–10^. Resident macrophages and DCs also produce cytokines and chemokines that help to recruit circulating neutrophils, monocytes, macrophages and DCs^11–13^. In mouse models of primary corneal HSV-1 infection, the timing and magnitude of these early innate responses are critical determinants of viral clearance and disease severity.

In addition to pathogen recognition receptors, cytokines and chemokines, HSV-1 infection is modulated by extracellular matrix components, but the underlying molecular mechanisms are ill-defined. Here we show that two corneal proteoglycans, lumican (Lum) and biglycan (Bgn) have significant roles in innate inflammatory responses and viral clearance in HSV-1 primary infections of the mouse cornea. Lum and Bgn are small leucine-rich repeat proteoglycans, abundant in the ECM of the corneal stroma where they bind with collagens and regulate collagen fibril architecture and organization ^14–16^. Our studies and those of others are uncovering a second role for these proteoglycans in cell communications in tissue injury, infection, inflammation and wound healing. Inflammatory signals upregulate expression of Lum ^17,18^ and Bgn ^19^ and even the epithelial layers induce their expression in the skin ^20^, gut ^17^ and in the cornea as we show here. Association with collagen in their ECM incorporated states, involves large segments of the Lum and Bgn core proteins ^21–24^, supporting the idea that when incorporated into stable ECMs they are less likely to be reactive with immune signals than the *de novo* synthesized forms ^18,25,26^. Consistent with this idea, collagen-free Lum upregulated in mouse models of corneal bacterial infection ^18^ and peritoneal sepsis promotes TLR4 responses ^27^. Bgn has been shown to promote both TLR2 and TLR4 signals in multiple inflammation settings ^19,28^. We discovered that both Lum and Bgn restrict TLR9 signals likely through direct binding with DNA ligands. The TLR9 regulations by Lum and Bgn are quite different from their signal-promoting activities of TLR2 and TLR4, which occur through direct interactions with TLR2/4 or the common partner CD14. As multiple TLRs are important in host response to HSV-1^4,29–31^, we sought to determine how Lum and Bgn regulate corneal response to HSV-1 infection. Using the HSV-1 keratitis mouse model, we show that mice lacking *Lum* or *Bgn* have poor viral clearance and develop more severe keratitis. We show blunted early induction of pro-inflammatory cytokines, chemokines, and type I IFNs, and poor early recruitment of monocytes and plasmacytoid dendritic cells (pDCs), compared to wild-type (WT) mice. To distinguish innate immune signals in cornea resident cells from those elicited in infiltrating immune cells we tested HSV-1 infection in stem cell-derived 3D human corneal organoids, which have a simple stratified epithelium and a LUM and BGN-rich corneal-like stroma. We found viral presence and induction of chemokines such as CCL2 in the epithelial layers. Together, our study indicates that host ECM proteins Lum and Bgn support early innate immune responses that may help to control viral infection and mitigate severe corneal inflammation.

## Results

### Increased disease score and poor viral clearance in *Lum^-/-^* and *Bgn^KO^* mice

We scratched and infected mouse corneas with 10^4^ pfu of HSV-1 (McKrae) in PBS and compared the response of the knockout strains with that of wild type (WT) mice. As controls, the corneas were scratched and coated with PBS alone. Clinical disease, assessed by slit-lamp biomicroscopy, show more pronounced corneal opacity and inflammatory changes in *Lum^-/-^* and *Bgn^KO^* mice compared to WT controls by day 3 post-infection (3 dpi). A blinded corneal scoring^32^ showed higher scores in the knockout strains compared to WT mice (**Fig. 1A-D**). Keratitis scores were significantly higher in *Lum^-/-^* (2.42 ± 0.38) and *Bgn^KO^* (1.43 ± 0.17) mice than in WT (1.27 ± 0.18) by 3 dpi. By 5 dpi, both knockout strains exhibited marked corneal clouding with dense central and sometimes pan-corneal haze, with the most severe phenotype noted in the *Lum^-/-^* mice (**Fig. 1C**). *Lum^-/-^* mice (2.44 ± 0.3) exhibited significantly higher keratitis scores than WT mice (0.57 ± 0.3), while clinical scores were also high in the *Bgn^KO^* mice (1.86 ± 0.51) these were not statistically significant (*p = 0.054*) (**Fig. 1D**). To evaluate whether enhanced disease severity in the knockout strains is due to impaired viral control, we quantified corneal viral titers by plaque assay at 3 and 5 dpi. At both time points, viral titers were higher in the *Lum^-/-^* (27 ± 3.27 and 102.5 ± 14.93 pfu, at 3 and 5 dpi respectively) and *Bgn^KO^* (26.17 ± 3.23 and 98 ± 16.55 pfu at 3 and 5 dpi) compared to WT mouse corneas (8 ± 2.76 and 15 ± 8.66 pfu at 3 and 5 dpi), demonstrating poor viral clearance in the absence of Lum and Bgn (**Fig. 1E**).

**Figure 1.**
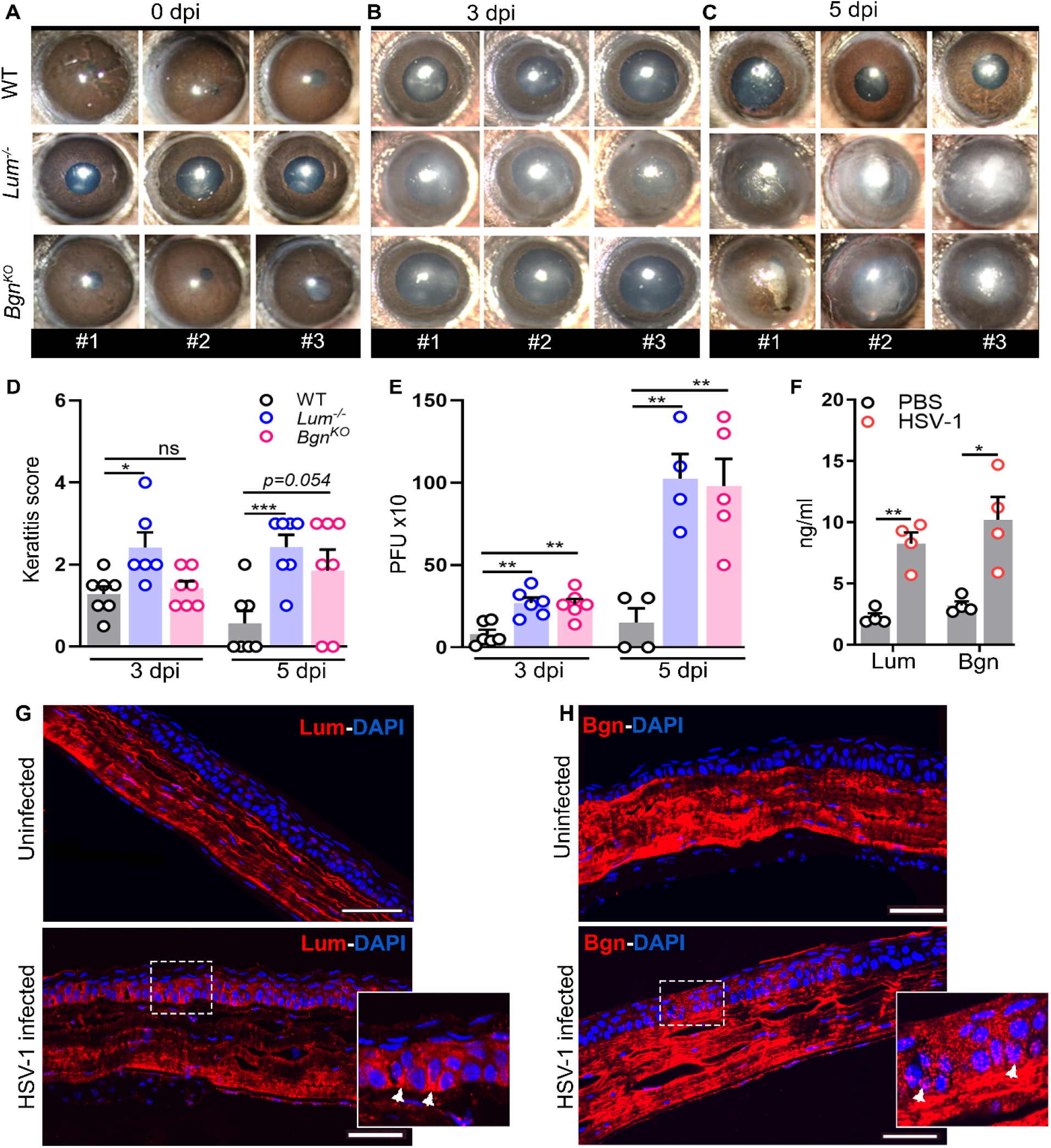
HSV-1-keratitis severity increases in *Lum*^-/-^ and *Bgn^KO^* mice. **(A–C)** Slit-lamp biomicroscopy images of corneas from three representative WT, *Lum*^-/-^, and *Bgn^KO^* mice shown from 3 independent experiments at **(A)** 0 days post-infection (dpi), **(B)** 3 dpi, and **(C)** 5 dpi with HSV-1. Progressive corneal opacity is evident in null mice compared to WT. **(D)** Cumulative keratitis scores at 3 and 5 dpi in WT, *Lum*^-/-^, and *Bgn^KO^* mice. n = 6 -7 mice/genotype from 3 independent experiments. *p < 0.05, ***p < 0.001; ns, not significant. **(E)** Viral titers in plaque-forming units (PFU) were measured from corneas of WT, *Lum*^-/-^, and *Bgn^KO^* mice at 3 and 5 dpi. n = 4 - 6 mice/genotype from 3 independent experiments. **p < 0.01. **(F)** ELISA based quantification of Lum and Bgn levels in corneal protein extracts from PBS-treated and HSV-1 infected mice. n = 4 mice/genotype from 2 independent experiments; *p < 0.05, **p < 0.01. **(G–H)** Immunohistochemical staining of corneal tissue sections from uninfected and HSV-1-infected mice at 5 dpi, demonstrating expression of **(G)** Lum and **(H)** Bgn by the basal epithelial cells during HSV-1 infection, which is absent in uninfected corneas. Insets show higher magnification; white arrowheads indicate epithelial cells expressing Lum and Bgn following infection. Scale bars, 100 µm. Image from one of 2 mice shown. Statistical significance for D, E and F was determined using two-way ANOVA with multiple comparisons analysis. Error bars represent the mean ± SEM.

Since both Lum and Bgn appear to be protective against HSV-1, we wondered whether their *de novo* expression increases during infection. We measured the levels of Lum and Bgn in the corneas of infected WT mice at 3 dpi by ELISA. Our data show that both Lum and Bgn proteins are significantly elevated in the HSV-1-infected corneal lysates compared to control PBS corneal lysates **(Fig. 1F)**. Immunolocalization of Lum and Bgn in WT corneas at 5 dpi show that beyond the bulk presence of these proteoglycans in the resting corneal stroma, in the infected corneas, they now appear in the basal epithelial cell layer as well (**Fig. 1G** and **H, Fig. S1A** and **B**). These findings suggest that Lum and Bgn are upregulated in the cornea, including the epithelium and contribute to early antiviral defense mechanisms in the cornea.

### Lum and Bgn support early innate immune cell recruitment to the infected corneas

We next questioned whether early immune cell recruitment was impaired in *Lum-* and *Bgn-*null mice. We used flow cytometry to immunophenotype the infiltrating leukocyte populations in WT, *Lum^-/-^*and *Bgn^KO^* mouse corneas following infection with HSV-1 at 3 and 5 dpi or PBS treatment as controls. Representative flow cytometric gating strategies for neutrophils (CD11b^+^Ly6G^+^), monocytes (CD11b^+^Ly6C^+^F4/80^-^), and inflammatory monocytes (CD11b^+^F4/80^+^Ly6C^+^) subsets are shown in **Fig. 2A**. The cumulative data (**Fig. 2B**) shows that at 3 dpi WT and *Bgn^KO^* mouse corneas had significantly higher numbers of neutrophil, 285 ± 40.13 and 381.6 ± 18.11 cells/cornea, respectively, compared to *Lum^-/-^* mouse corneas which on average displayed 74.2 ± 7.39 cells/cornea. By 5 dpi the neutrophil numbers in WT were back to basal levels, while these were elevated in *Lum^-/-^* and *Bgn^KO^*corneas without reaching statistical significance. The monocyte population showed a similar trend at 3 dpi; large numbers in WT (479.2 ± 26.52 cells/cornea) and significantly lower numbers in *Lum^-/-^* (90.2 ± 6.97 cells/cornea) and *Bgn^KO^* (293 ± 40 cells/cornea) (**Fig. 2C)**. By 5 dpi monocyte numbers in WTs remained almost as high as their levels at 3 dpi, while the numbers showed some increase in *Lum^-/-^* and *Bgn^KO^* corneas. When we examined the inflammatory monocyte subset CD11b^+^F4/80⁺Ly6C^hi^), while the absolute numbers were low (< 100 cells/cornea) in all strains, the *Lum^-/-^* corneas showed significantly lower than WT numbers at 3 dpi (**Fig. S2**).

**Figure 2.**
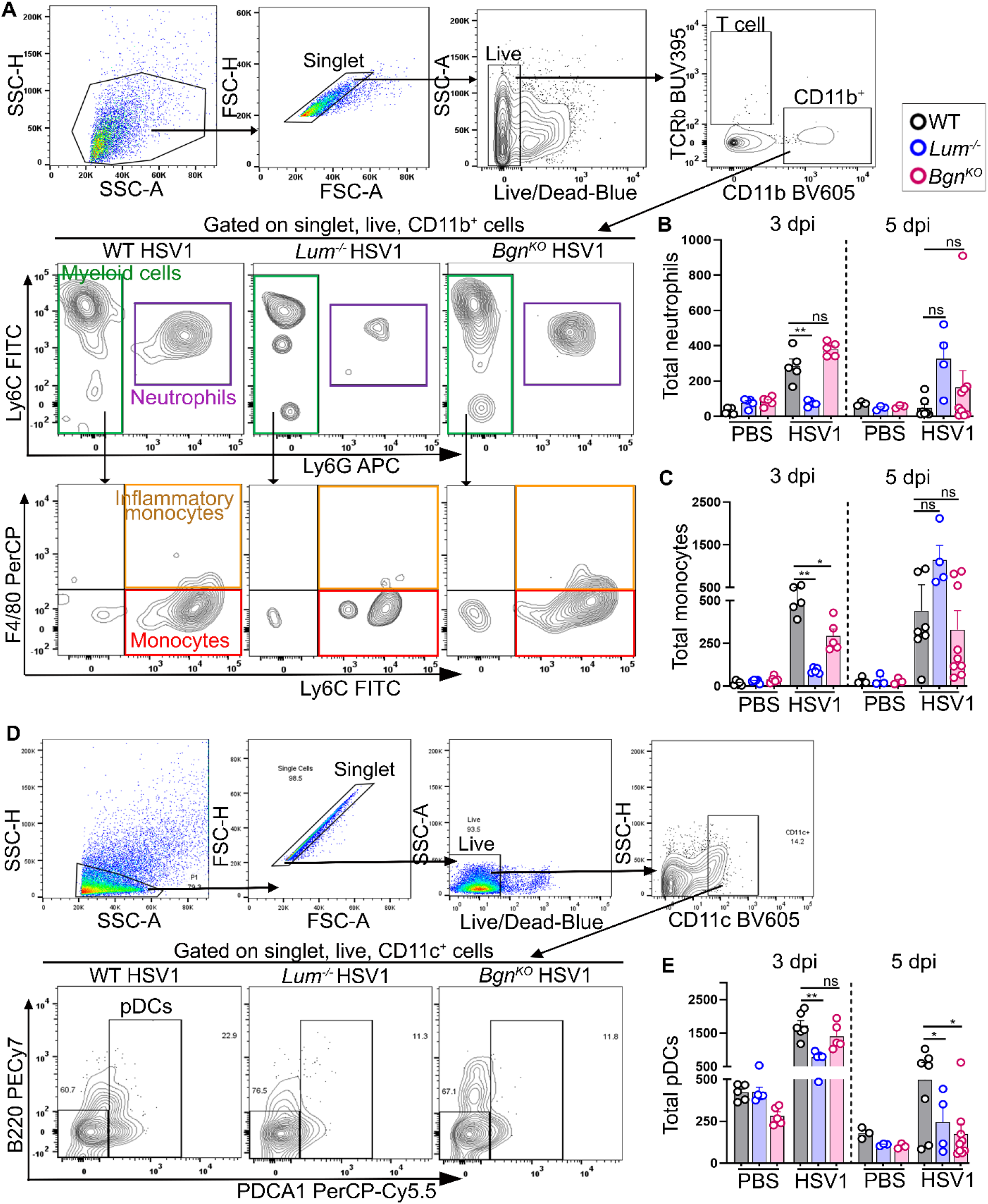
Poor immune cell infiltration in HSV-1 infected *Lum*^-/-^ and *Bgn^KO^* mouse corneas. **(A)** Representative gating strategy for monocytes (Ly6C⁺F4/80^-^Ly6G⁻), neutrophils (Ly6C^-^ Ly6G^+^) and inflammatory monocytes (Ly6C⁺F4/80^+^Ly6G⁻) in the cornea. Cells were gated on singlets, live cells and CD11b⁺ cells (TCRβ⁻ CD11b⁺). Representative contour plots are shown for HSV-1 infected WT, *Lum*^-/-^, and *Bgn^KO^* mice **(B)** Quantification of total neutrophils in corneas. n = 3 – 9 mice/genotype; **p < 0.01; ns, not significant. **(C)** Quantification of total monocytes in corneas. n = 3 – 9 mice/genotype; *p < 0.05, **p < 0.01; ns, not significant. **(D)** Representative gating strategy for pDCs (CD11c^+^B220⁺ PDCA1⁺) in corneal tissue. Cells were gated on singlets, live cells, and CD11c⁺ cells. Representative contour plots are shown for HSV-1 infected WT, *Lum*^-/-^, and *Bgn^KO^* mice. **(E)** Quantification of total pDCs in corneas. Error bars represent mean ± SEM, n = 3 – 9 mice/genotype; *p < 0.05, **p < 0.01; ns, not significant. Statistical significance for B, C and E was determined using unpaired t-test with Mann-Whitney correction. Error bars represent the mean ± SEM from 3 independent experiments.

Both resident and recruited plasmacytoid dendritic cells (pDC) are crucial for mounting an antiviral response against HSV-1 ^33,34^. We found that pDC numbers were significantly lower in *Lum^-/-^* corneas compared to WT at 3 dpi, while the numbers in *Bgn^KO^* corneas were like WTs. By 5 dpi pDC numbers were low in both knockout genotypes 174 ± 60 cells/ *Lum^-/-^* cornea and 74 ± 7 cells/ *Bgn^KO^* cornea) compared to WTs (497 ± 128 cells/cornea) (**Fig. 2D** and **E**). Not surprisingly, at these early time points, the CD4+ and CD8+ T cell populations in the infected corneas were low and similar across all three genotypes (**Fig. S3A – C**). We also assessed the numbers of myeloid and lymphoid cells in the draining lymph node (dLN) of infected and PBS-treated control mice. Total neutrophil, monocyte, pDC, CD4+, and CD8+ T cell numbers in the dLN of infected cornea did not differ significantly among genotypes at 3 dpi (**Fig. S4A**). However, at 5 dpi, total neutrophils and pDCs in the dLN of infected cornea were significantly higher in the WT (1197 ± 19 cells/dLN) compared to *Lum^-/-^* (450 ± 18 cells/dLN) and *Bgn^KO^* (600 ± 13 cells/dLN) (**Fig. S4B**). Altogether, these data indicate that Lum and Bgn are required for optimal early recruitment of inflammatory monocytes and plasmacytoid DCs into the HSV-1– infected cornea, whereas neutrophil infiltration was most impaired by the absence of Lum.

### Lum and Bgn deficiency impairs the innate immune response in HSV-1 infection

Given the lower recruitment of immune inflammatory cells in the infected null mouse corneas, we performed bulk RNA sequencing of infected and uninfected corneas at 5 dpi to assess how innate inflammatory genes are regulated in the WT and the null mutant corneas. Principal component (PC) analysis showed clear segregation of infected corneal expression from the PBS-treated controls along the first two principal components, indicating that as one would expect that infection causes a fundamental change in gene expression (**Fig. 3A**). Volcano plots show that genes most differentially expressed in all infected genotypes compared to their strain-specific uninfected control corneas were generally similar. Type I IFN pathway genes and interferon-stimulated genes (ISGs) and canonical antiviral mediators such as *Mx1*, *Slfn4, Rsad2, Gbp5, Mnda* and *Oas1g* were stimulated in all. In addition, the chemokines Ccl2 and Cxcl10 were also upregulated in the corneas of all three infected strains (**Fig. 3B – D**). Gene Ontology (GO) enrichment analysis of biological processes was performed on the upregulated DEGs in HSV-1-infected corneas (padj ≤ 0.05, log2FC > 1.5). In all three genotypes, the enriched biological processes were dominated by antiviral and immune response categories, including regulation of immune effector processes and cytokine-mediated signaling pathways and leukocyte-mediated immunity. We also noted that leukocyte-mediated immunity and lymphocyte activation were the most significant groups in the WT and *Bgn*^KO^ mice **(Fig. 3E** and **G)**. Thus, bulk transcriptomic data at 5 dpi suggested upregulations in innate inflammatory signaling across all genotypes. However, the flow cytometry-based immune cell phenotyping presented earlier indicated clear differences in innate immune cell activities as early as 3 dpi, with *Lum^-/-^* corneas, in particular showing cytokine and chemokine levels lower than those of WT. This suggested that transcriptomic similarities at 5 dpi mask functional deficits in early innate immune responses that likely differ between the WT and mutant strains.

**Figure 3.**
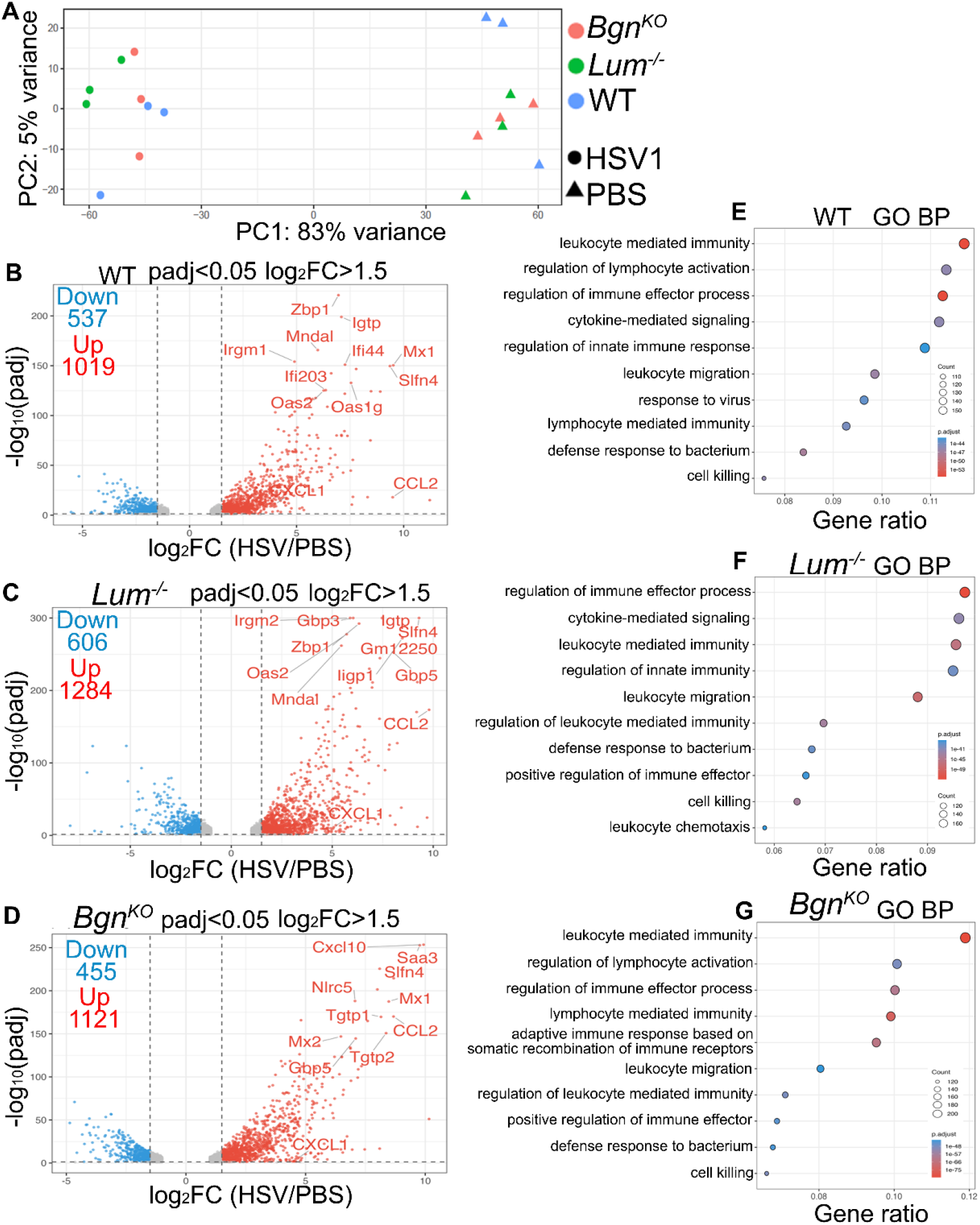
Bulk RNA-seq showing upregulation of inflammatory and interferon-stimulated genes in HSV-1 infected corneas. **(A)** Principal component analysis of bulk RNA-seq data from corneas of WT, *Lum*^-/-^, and *Bgn^KO^* mice infected with HSV-1 or PBS control. PC1 and PC2 account for 83% and 5% of total variance, respectively, demonstrating clear separation between HSV-1-infected and PBS-treated groups across all genotypes. **(B–D)** Volcano plots showing DEGs in HSV-1 infected versus PBS-treated control corneas for **(B)** WT, **(C)** *Lum*^-/-^ and **(D)** *Bgn^KO^* mice. Significantly upregulated and downregulated genes (padj < 0.05, log₂FC > 1.5) are shown in red and blue, respectively; non-significant genes are shown in gray. Selected top differentially expressed genes are labeled, including chemokines such as *CCL2* and CXCL1 and interferon-stimulated genes such as *Mx1*, *Slfn4*, *Oas2*, and *Zbp1*. **(E–G)** Gene Ontology Biological Process (GO BP) enrichment dot plots for significantly upregulated genes in HSV-1-infected **(E)** WT, **(F)** *Lum*^-/-^ and **(G)** *Bgn^KO^* mice corneas. Enriched GO BP include leukocyte-mediated immunity, cytokine-mediated signaling, regulation of innate immune response and leukocyte migration. Of note, leukocyte-mediated immunity was the most significant group in the WT and *Bgn*^KO^ mice.

We posited that the null mutant strains may have delayed and reduced induction of proinflammatory cytokines and chemokines early after HSV-1 exposure. To address this question, we evaluated selected proinflammatory cytokines and chemokines in the cornea using multiplex cytokine assays (LEGENDplex, BioLegend) at 3 and 5 dpi. We observed that while the pro-inflammatory cytokines, TNF-α, IFN-γ, IL12p70, IL-6, IL-1β and GM-CSF were clearly induced in all three strains, they were significantly lower in the infected null compared to WT corneas. By 5 dpi all of these pro-inflammatory cytokines were reduced to almost basal levels in the WT infected corneas (**Fig. 4A**) The chemokines CXCL1, CXL10, CCL2 and CCL5, were also induced in all three genotypes compared to their uninfected control corneas, but the levels were two to three times higher in WT than *Lum^-/-^* or *Bgn^KO^* mouse corneas (**Fig. 4B**). Similarly, at 3 dpi the anti-inflammatory cytokine IL-10 was induced in all three genotypes, but the increase in WT mice corneas were twice as much as that seen in the null mouse corneas (**Fig. S5**). At 3 dpi, levels of type I interferons, IFN-α and IFN-β, were significantly lower in the knockouts compared to WT. At 5 dpi, IFN-α levels showed a modest increase in the infected knockout corneas, while it was almost reduced to basal levels in the WT and IFN-β levels were low in all infected corneas (**Fig. 4C**).

**Figure 4.**
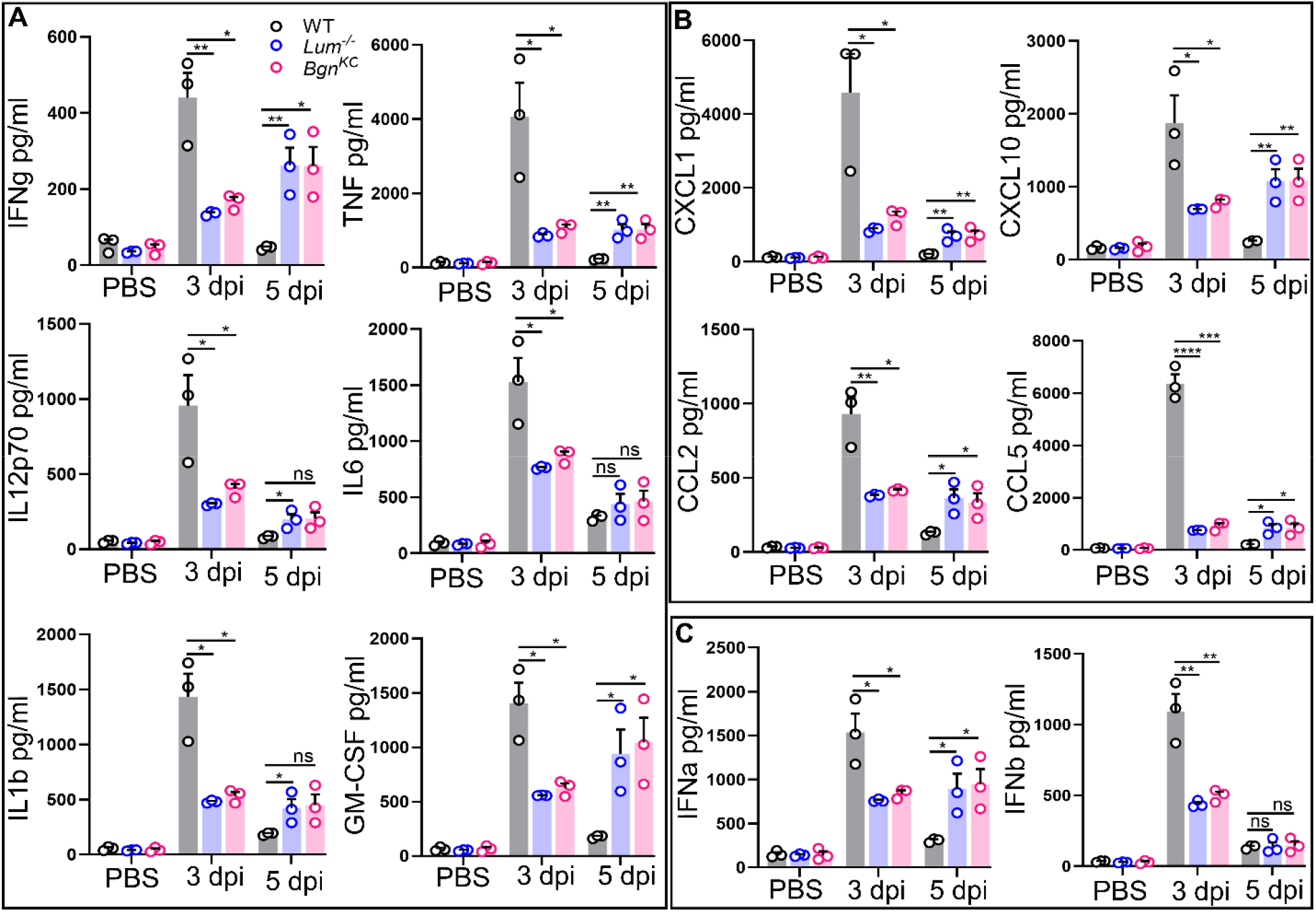
Lum and Bgn deficiency attenuates cytokine and chemokine production in the HSV-1 infected corneas. **(A)** At 3 dpi corneal homogenates from WT mice show robust induction of pro-inflammatory cytokine levels including IFN-γ, TNF, IL-12p70, IL-6, IL-1β, and GM-CSF, while *Lum*^-/-^ and *Bgn^KO^*show much lower levels of induction. By 5 dpi, cytokine levels declined in WT mice, while *Lum*^-/-^ and *Bgn^KO^* corneas show small increases. n = 3 mice/genotype; *p < 0.05, **p < 0.01; ns, not significant. **(B)** At 3 dpi corneal homogenates from WT mice show significant increases in CXCL1, CXCL10, CCL2, and CCL5, while *Lum*^-/-^ and *Bgn^KO^* mouse corneas show some increases over their PBS-controls, these were much lower than the inductions in WT. By 5 dpi, cytokine levels declined in WT mice, while *Lum*^-/-^ and *Bgn^KO^* mice showed increased levels of chemokines. n = 3 mice/genotype; *p < 0.05, **p < 0.01; ***p < 0.001, ****p < 0.0001, ns, not significant. **(C)** Type I interferon levels (IFN-α and IFN-β) in HSV-1 infected WT mice corneal homogenates were significantly higher At 3 dpi compared to *Lum*^-/-^ and *Bgn^KO^* mice, with no significant difference in IFN-β observed at 5 dpi among genotypes. Error bars represent mean ± SEM, n = 3 mice/genotype; *p < 0.05, **p < 0.01; ns, not significant. Statistical significance for A, B and C was determined using an unpaired t-test with Mann-Whitney correction. Error bars represent the mean ± SEM.

Taken together, the results show that WT corneas display sharp early (3 dpi) antiviral responses, while *Lum^-/-^* and *Bgn^KO^* mice show a delayed (5 dpi) induction that fails to reach WT levels. Our data suggest that loss of Lum and Bgn results in delayed and substantially lower early production of antiviral cytokines, chemokines, and type I IFNs in the cornea, consistent with lower than WT recruitment of innate immune cells, poor viral clearance, and increased corneal disease scores.

### Early Ccl2 induction in WT mice may have corneal epithelial cell contributions

As Ccl2, a potent chemoattractant for monocytes^35^, was highly induced in WT mouse corneas by 3 dpi, we wondered whether some of this early Ccl2 response was from the infected corneal epithelial layers. To test this, we performed immunohistochemical staining of corneal sections from WT, *Lum^-/-^* and *Bgn^KO^* mice following HSV-1 infection. By 3 dpi, WT corneas showed strong Ccl2 immunoreactivity that was predominantly localized to the corneal epithelium with scattered signal within the stromal cells (**Fig. 5A**). In contrast, the HSV-1 infected *Lum^-/-^* and *Bgn^KO^* corneas had significantly lower Ccl2^+^ cells in the epithelium compared to WT (**Fig. 5B**). The uninfected WT mice corneas lacked Ccl2+ cells, and the no primary control (NPC) confirmed the specificity of anti-Ccl2 primary antibody (**Fig. S6A - B**). These findings identify the corneal epithelium as a major source of monocyte-recruiting signals during early HSV-1 infection and suggest that Lum and Bgn facilitate epithelial Ccl2 production *in vivo*. Immunolocalization data presented earlier show that Lum and Bgn are transiently induced in the epithelial layers following HSV-1 infection and may act in an autocrine manner to facilitate epithelial chemokine production during the critical early phase of the innate immune response.

**Figure 5.**
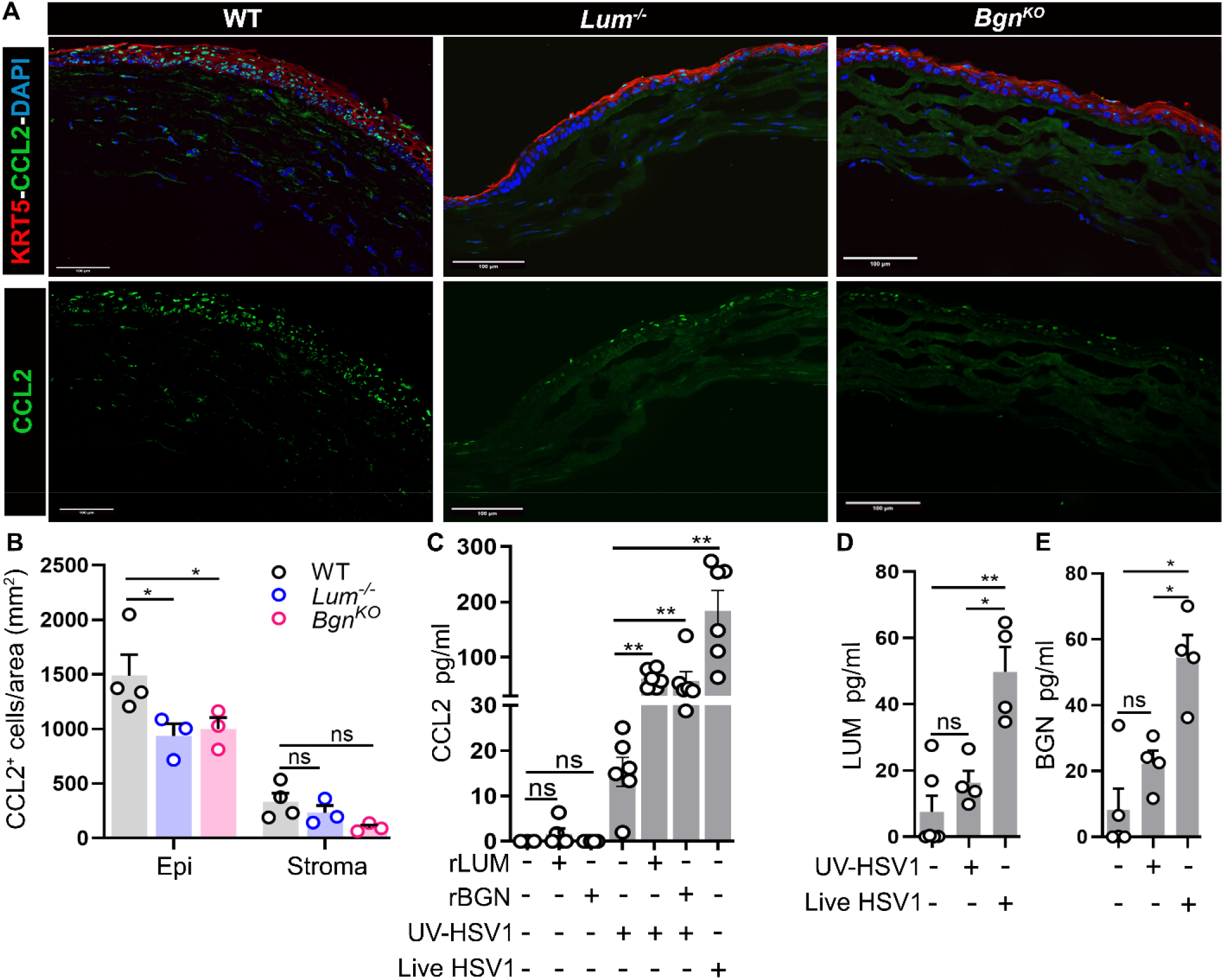
Lum and Bgn promote Ccl2 expression in the corneal epithelium during HSV-1 infection. **(A)** Representative immunohistochemical staining of corneal sections from HSV-1-infected WT, *Lum*^-/-^ and *Bgn^KO^* mice at 3 dpi demonstrates strong Ccl2 immunoreactivity throughout the epithelium and stroma in WT corneas, and markedly reduced staining in *Lum*^-/-^ and *Bgn^KO^* mouse corneas. Scale bars: 100 µm. One representative image from 3 mice/genotype. **(B)** Quantification of Ccl2⁺ cell numbers from sections of 3 dpi WT, *Lum*^-/-^ and *Bgn^KO^* corneas. WT mice showed significantly more Ccl2⁺ corneal epithelial cells than both null strains. No significant differences were detected in the stromal compartment. *p < 0.05; ns, not significant. **(C)** Secreted CCL2 (pg/mL) measured by ELISA in supernatants from hTCEpi cells show that recombinant lumican (rLUM) or rBGN upregulate CCL2 in cells treated with UV-inactivated HSV-1 (UV-HSV1). Neither rLUM or rBGN alone stimulate CCL2 production. Cells infected with live HSV-1 alone used as a positive control show maximal CCL2 production. **p < 0.01; ns, not significant. **(D–E)** ELISA quantification of secreted endogenous **(D)** LUM and **(E)** BGN show their significant upregulation in response to the live HSV-1 infection and not UV-HSV1 or untreated controls. *p < 0.05, **p < 0.01; ns, not significant. Statistical significance for B, C, D and E was determined using an unpaired t-test with Mann-Whitney correction. Error bars represent the mean ± SEM.

#### LUM and BGN promote HSV-1-triggered CCL2 induction in human corneal epithelial cells *in vitro*

We next investigated whether LUM and BGN can modulate epithelial chemokine responses *in vitro* using the human telomerase-immortalized corneal epithelial (hTCEpi) cell line. The hTCEpi cells were exposed to UV-inactivated HSV-1 in the presence or absence of recombinant human lumican (rLUM) or biglycan (rBGN). UV-inactivated HSV-1 (McKrae) infection (MOI-1) alone triggered robust CCL2 protein secretion compared to uninfected hTCEpi controls, indicating that corneal epithelial cells can directly sense HSV-1 components and mount a chemokine response. Remarkably, the addition of human rLUM or rBGN significantly enhanced CCL2 production compared to UV-inactivated HSV-1 (UV-HSV-1) infection alone (**Fig. 5C & Fig. S6C)**. Neither rLUM nor rBGN alone induces CCL2 in the absence of HSV-1, indicating that these proteoglycans are not primary chemokine inducers but enhance virus-triggered epithelial signaling. Also, the UV-HSV-1 infected and uninfected hTCEpi cells do not secrete significant levels of LUM or BGN, but live HSV-1 (MOI-1) induces significant LUM and BGN secretion into the culture media (**Fig. 5D – E**). Altogether, these results demonstrate that LUM and BGN are induced in the epithelial layers after HSV-1 infection and they enhance CCL2 production by corneal epithelial cells, both in vivo and in cell culture. By promoting virus-induced epithelial signaling and CCL2 secretion, Lum and Bgn promote efficient recruitment of neutrophils and monocytes to the infected cornea.

### Human corneal organoids recapitulate HSV-1-driven epithelial CCL2 induction

To validate epithelial CCL2 induction upon HSV-1 infection in physiologically relevant tissue architecture, we leveraged our well-characterized human iPSC-derived corneal organoid (HCO) model, which recapitulates native corneal structure ^20,36,37^. Our previous scRNA-seq study showed that the HCOs exhibit key features of the native cornea, including a stratified epithelium and an organized stromal compartment, and express many early innate immune sensors for microbial components. HCOs were infected with 1000 pfu of the mCherry-expressing HSV-1 KOS strain^38^ and analyzed 24 hours post-infection (hpi). The infected HCOs displayed discrete, bright mCherry^+^ foci on the organoid surface, indicating robust viral adherence and epithelial tropism (**Fig. 6A**). Immunohistochemical analysis of uninfected HCOs revealed well-organized KRT3^+^ stratified epithelial layers overlying a COL5A1^+^ stromal compartment **(Fig. 6B)**. Notably, HSV-1-infected HCOs showed viral signal predominantly within the KRT3^+^ epithelial layers, closely recapitulating the epithelial tropism characteristic of native corneal HSV-1 infection **(Fig. 6C)**. HCO cryosections immunostained for CCL2 showed intense cytoplasmic immunostaining of epithelial cells within the cornea-like basal epithelial layer. No CCL2 was detected in immunostaining of uninfected HCOs (not shown). Co-staining with KRT3 confirmed that the CCL2-producing cells were corneal epithelial, whereas the underlying stromal-like regions showed a comparatively weak signal **(Fig. 6D – E)**. We detected increased CCL2 levels by ELISA in the tissue culture media of infected HCOs **(Fig. 6F)**. Taken together, the HCO model recapitulates the key features of epithelial Ccl2 induction we observed in infected mouse corneas and cultured human corneal epithelial cells. Most notably, HSV-1 infection, epithelial tropism of the virus and the epithelial induction of CCL2 raises the HCO as a suitable corneal model.

**Figure 6.**
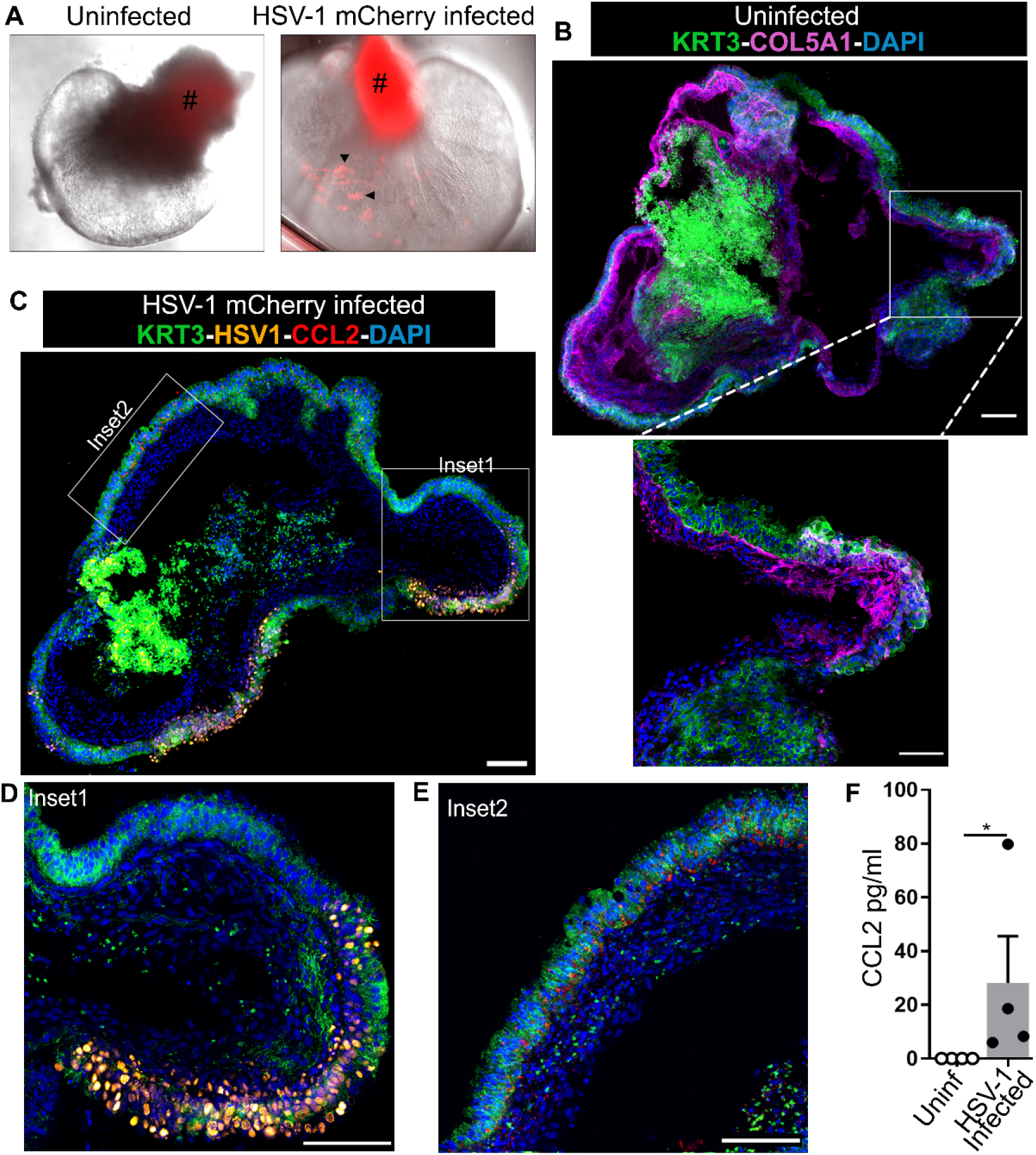
CCL2 is induced in the epithelial layers of HSV-1 Infected human corneal organoids (HCO). **(A)** Representative brightfield images of uninfected HCO and 24 hpi with HSV-1 expressing mCherry (red). Arrowheads indicate foci of mCherry^+^HSV-1 infection visible at the organoid periphery, confirming successful viral infection. (#) denotes autofluorescence inn the pigmented HCO stalks. **(B)** Representative image shows typical immunohistochemical staining of corneal epithelial marker KRT3 and stromal marker COL5A1 in uninfected HCO at 20× magnification. The inset below shows 40x magnification view of the organized layering of KRT3⁺ epithelial cells, with a COL5A1⁺ stromal layer below. Scale bars: 100 µm. **(C)** Representative image showing HSV-1 mCherry-infected HCO. HSV-1 infection is localized to discrete foci within the KRT3⁺ epithelial layer. CCL2 expression is detected in the epithelial compartment of infected HCOs. Scale bar: 100 µm. **(D-E)** Higher-magnification view of Inset 1 from panel C, showing HSV-1 (orange) within KRT3⁺ (green) epithelial cells surrounding a region of active viral replication **(D)**. Inset 2 from panel C, depicting the peripheral epithelial region of the HCO with uninfected bystander KRT3⁺ epithelial cells expressing CCL2⁺ (red) in areas distal to the primary infection focus **(E)**. Scale bar: 100 µm. **(F)** ELISA quantification of CCL2 protein levels in supernatants collected from uninfected (Uninf) and HSV-1-infected HCOs demonstrates CCL2 secretion from the infected HCOs. Statistical analysis was done using an Unpaired t-test with Mann-Whitney correction. Error bars represent mean ± SEM; *p < 0.05.

## Discussion

Here we show that mice lacking lumican or biglycan have worse HSV-1 mediated corneal disease. We observed a clear decrease in viral clearance in nulls as early as 3 dpi and higher clinical corneal disease scores, compared to WT mice. We found a significant impairment in neutrophil and monocyte infiltration in *Lum^-/-^* and *Bgn^KO^* and worse clinical disease compared to WT. Of relevance to our findings, a previous study reported the early homing of neutrophils, and monocytes in particular to the corneal epithelium as having a protective role against viral dissemination^39^. Another study reported that neutrophil-mediated NETosis can restrict HSV-1 replication and viral dissemination in the mouse cornea^40^. In addition to neutrophils and monocytes, pDCs have a significant role in HSV-1 control in the cornea. The pDCs are not only the major source of type-I IFNs, but they also produce multiple cytokines (IL-12p70, TNF, IFNγ and IL-6) and chemokines (CCL3 and CXCL10) that orchestrate the activation of other innate immune cells like NK cells during viral infection^34,41^. We found infiltration of pDCs were also lower in *Lum^-/-^* and *Bgn^KO^*at these early stages of infection. In the future, it will be interesting to investigate the role of Lum and Bgn in regulating the pDC-NK cell axis. In step with reduced or delayed immune cell infiltration, we observed less than WT levels of pro-inflammatory (IFN-γ, IL-1b, TNF, GM-CSF and IL-12) cytokine and chemokine (CCL2, CCL5, CXCL1 and CXCL10) induction in infected *Lum^-/-^* and *Bgn^KO^* corneas. We also noted that immune-protective IL-10 was not induced well in the HSV-1-infected null-mouse corneas compared to WTs. Ocular IL-10 is known to reduce HSV-1-infected corneal disease by suppressing overactive pro-inflammatory cytokine and chemokine secretion^42^. That Lum and Bgn actively regulate HSV-1 infection is further supported by our observations that, at the protein level, both are elevated in the corneas of WT infected mice at 3 dpi. In cultured epithelial cells infected with the live virus, we also found increased secretion of Lum and Bgn in the media. While no other studies have examined the role of lumican or biglycan in HSV-1 ocular infections, one study on intervertebral disc degeneration reported decreased biglycan transcript in human nucleus pulposus cells after HSV-1 infection in culture^43^. These differences in expression in experimental settings likely arise due to differences in cell types and protein level (in our study) versus transcript level measurements. Both Lum and Bgn are detectable and often elevated in circulation in disease settings ^25,44–46^.

Although we found absolute lymphocyte numbers to be low as is expected at these early time points, Lum and Bgn may shape adaptive immune responses in later phases of infection. Reduced early recruitment of monocytes and pDCs, together with attenuated cytokine and chemokine production in *Lum^-/-^* and *Bgn^KO^* corneas, may influence subsequent T cell priming, trafficking, and effector function. This possibility is consistent with our previous findings in allergic contact dermatitis, where we found T cell-mediated inflammatory responses to be modulated by Lum and Bgn, supporting a broader role for these ECM proteoglycans in linking innate and adaptive immunity^47^.

Our bulk RNA-seq at 5 dpi showed that WT, *Lum^-/-^* and *Bgn^KO^* corneas all upregulated broad antiviral and immune-response programs, including type I IFN pathway genes, ISGs, and chemokines such as *Ccl2* and *Cxcl10*. By contrast, multiplex cytokine analysis revealed a marked deficit in the early inflammatory response in the null mouse corneas at 3 dpi, including reduced levels of proinflammatory cytokines, chemokines and type I IFNs relative to WT. By 5 dpi, some antiviral and inflammatory transcripts were induced across all genotypes, suggesting that the knockout corneas eventually mount a partial response. However, this delayed transcriptional activation does not compensate for the early deficit in cytokine and chemokine production, which coincides with reduced recruitment of neutrophils, monocytes, and pDCs and with impaired viral clearance. These findings suggest that Lum and Bgn likely support timely amplification of the early inflammatory signals that coordinate effective innate control of HSV-1 in the cornea.

The interactions of Lum and Bgn with inflammatory signals may be largely dominated by their known cross talks with TLR2, 4 and 9. We uncovered a novel TLR 9 regulation by Lum and Bgn; both bind CpG DNA and limit TLR9 response in cell culture ^27^. In a mouse sepsis model, we further found that compared to wild types, the *Lum*^-/-^ displayed lower TLR9 response (phosphorylation of IRF7, lower type I IFN) suggesting that Lum limits TLR9 signals in vivo. In addition, we detected elevated circulating Lum and Bgn in human sepsis patients. On the other hand, both Lum and Bgn increase TLR4 signals in cell culture and mouse models ^17–19,27,48–50^. Previously, we found that *Lum^-/-^*showed increased mortality in live *Pseudomonas aeruginosa* lung infections ^27,49^ and in corneal infections, poor bacterial control and worse disease ^18^. The *Bgn^KO^* mice were reported to be partially resistant to experimental LPS-sepsis ^19,50^. We found that Lum was able to bind CD14 and Caveolin 1 and increased TLR4 residence on the plasma membrane of macrophages, while endocytosed Lum delayed the lysosomal degradation of TLR4. Bgn promotes TLR2 and TLR4 signals, presumably through CD14. Thus, the dampened early inflammatory signals in the HSV-1 infected null mouse corneas may be due to the lack of Lum or Bgn mediated TLR 2 and 4 signal enhancements. On the other hand, TLR9 signaling is increased in the absence of Lum and Bgn, and this may cause per cell increased type I interferon induction in pDCs of the null mice. But overall low infiltration of pDCs due to poor innate immune signals in the null corneas ultimately yield lower total type I interferons and antiviral protection. Another consideration is that the TLR9-restrictive functions of Lum and Bgn in WT corneas may help to control excessive TLR9 activation and thereby limit type I IFN-mediated tissue damage^51,52^.

Lum and Bgn, have other interactions with immune signals that are not fully understood that may be relevant to early innate defense. While preliminary observations (our unpublished data) show no interactions between Lum or Bgn and cGAS-STING, their DNA -binding properties may have a role in masking illicit microbial or self-DNA recognition by cGAS. Lum also interacts with CD18/β2 integrin that supports neutrophil migration ^53^, its interactions with CD14 and CD18 may also regulate phagocytosis which may modulate viral and dead cell clearance^49,54^.

Our findings that Lum and Bgn regulate Ccl2 production in epithelial layers is entirely novel and may have a significant early protective role against HSV-1 in mice and human primary infections. Ccl2 is induced by activated NFκB downstream of many different innate immune signals ^55,56^. Infected WT corneal epithelial layers showed marked immunostaining of Lum and Bgn, as well as Ccl2. Treatment of a human epithelial cell line with exogenous rLUM or rBGN upregulated CCL2 release upon treatment with HSV-1, while viral encounter itself also upregulated endogenous LUM and BGN secretion in culture. Our 3-dimensional human corneal organoids which have some of the architecture of the corneal epithelium and stroma without resident or infiltrating immune cells, also showed clear CCL2 immunostaining of epithelial layers adjacent to visibly infected areas. A recent study reported that Ccl2 produced by corneal sensory neurons have an anti-viral role in HSV-1 corneal infection^57^. Our findings here that an epithelial Lum/Bgn – Ccl2 axis may well add to this Ccl2-mediated anti-viral defense.

### Limitations of the study

There are several limitations in the current study that we should keep in mind. First, to tease out the intricate interplay between Lum and Bgn and the different TLRs in infection settings, further study of tissue-specific *Lum*- and *Bgn*-deficient mice and *Tlr*-knock out mice are needed. Our newly found regulations of epithelial Ccl2-induction require additional studies using conditional epithelial deletions of *Lum* and *Bgn* in mice to decipher contributions of epithelial versus Lum and Bgn released by a remodeling ECM stroma. Our multiplex cytokine analysis at 3 dpi revealed a striking early defect in innate immune response in *Lum^-/-^* and *Bgn^KO^* corneas compared to WTs. Yet, our bulk RNA-seq at 5 dpi did not discern major transcriptional differences in innate immune programs between the null and WT infected corneas. This suggests a complex transient early interaction between Lum/Bgn and host innate immunity that is important. Finally, bulk transcriptomic profiling is inherently limited in resolving cell-type-specific responses, highlighting the need for greater resource-dependent single-cell approaches to define the Lum and Bgn-mediated cellular drivers of antiviral immunity.

In summary, we found a key protective role for Lum and Bgn in HSV-1 infections of the cornea that likely engage their TLR2 and 4 signal-promoting functions. We further detected HSV-1-mediated induction of Lum and Bgn in the corneal epithelium and their direct contributions in elevating release of the CCL2 chemokine.

## Acknowledgments

We thank Drs Prashant J. Desai, The Johns Hopkins University School of Medicine and Daniel J. Carr, University of Oklahoma for sharing the HSV-1 mCherry (KOS strain) and HSV-1 (McKrae), respectively. We thank the NYU Langone Microscopy Laboratory (RRID: SCR_017934), Experimental Pathology Research Laboratory (RRID: SCR_017928) and Cytometry and Cell Sorting Laboratory which are partially supported by the Laura and Isaac Perlmutter Cancer Center support grant P30CA016087 and the Shared Instrument Grant S10 OD021747. We thank Michael Cammer for advice on confocal microscopy.

## Author contributions

Conceptualization: GM, SC; Methodology: GM, MK, ES, TS; Investigation: GM, MK, ES; Bioinformatics data analysis: YS, AKJ, GM, SC; Validation: GM, SC; Data Analysis: GM, MK, HY, AJSL, SC; Supervision: SC, GM; Writing—original draft: GM and SC; Review & editing: GM, MK, ES, YS, TS, AKJ, HY, AJSL, SC; Funding: GM, SC.

## Lead Contact

Requests for further information and resources should be directed to and will be fulfilled by the lead contact, Shukti Chakravarti.

## Declaration of generative AI in scientific writing

During the preparation of this work, the authors used ChatGPT (OpenAI) to assist with improving grammar and clarity of the text. After using this tool, the authors reviewed and edited the content as needed and take full responsibility for the content of this article.

## Data and materials availability

This study did not generate new unique reagents. All data needed to evaluate the conclusion in the paper are available in the main text and the supplementary materials. All raw bulk RNA-seq datasets will be deposited and made publicly available through the GEO database.

## Supplemental information

Supplemental Figures S1 - S6.

## Key resources table

*The key resources table (KRT) serves to highlight materials and resources essential to reproduce results presented in the manuscript. The items in the table must also be reported alongside the description of their use in the method details section. Literature cited within the KRT must be included in the references list. Please do not add custom headings or subheadings to the KRT. We highly recommend using RRIDs (see* https://scicrunch.org/resources*) as the identifier for antibodies and model organisms in the KRT. To create the KRT, please use the template below or the <u>KRT webform</u>. See the more detailed <u>Word table template</u> document for examples of how to list items*.

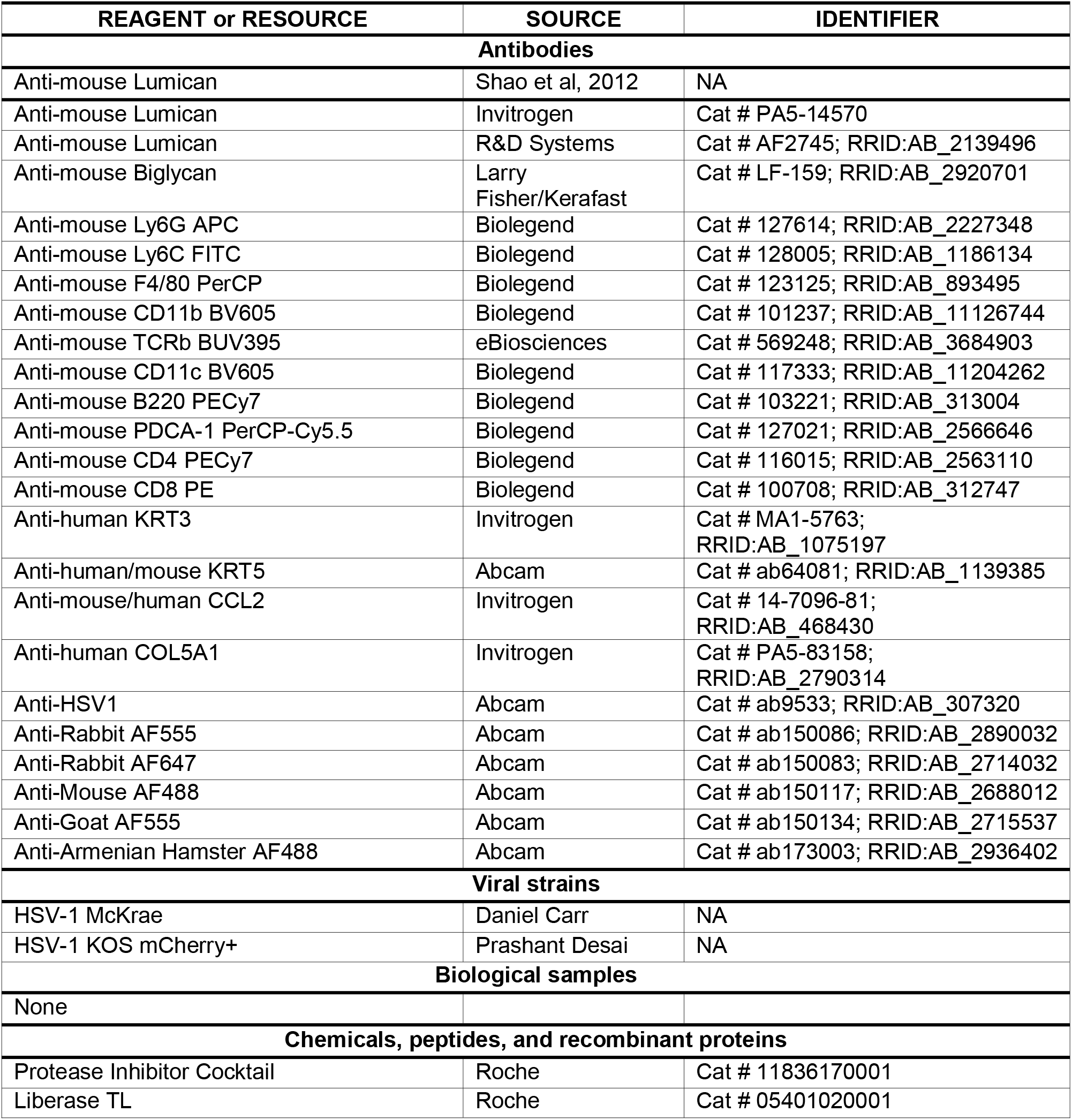
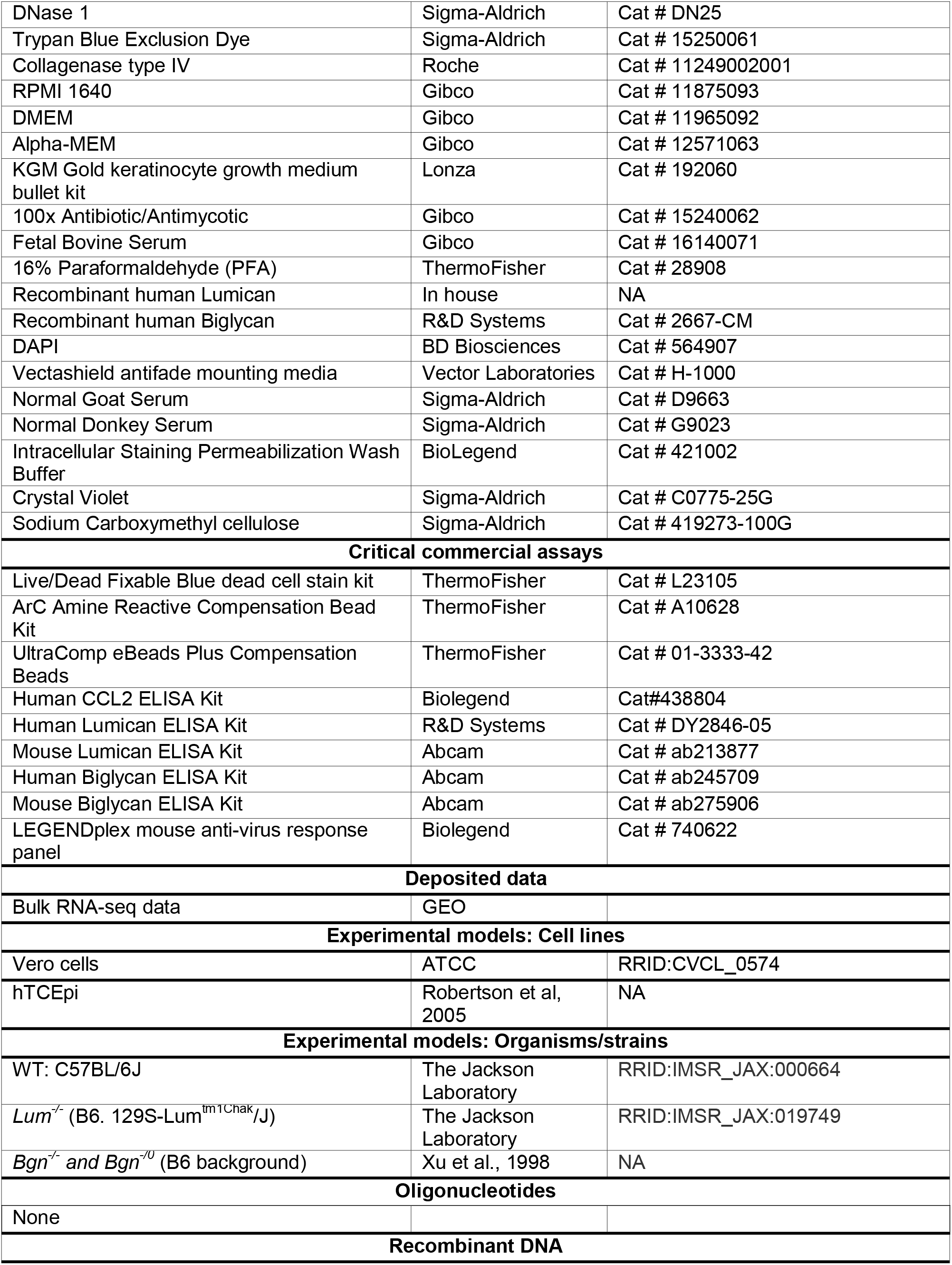
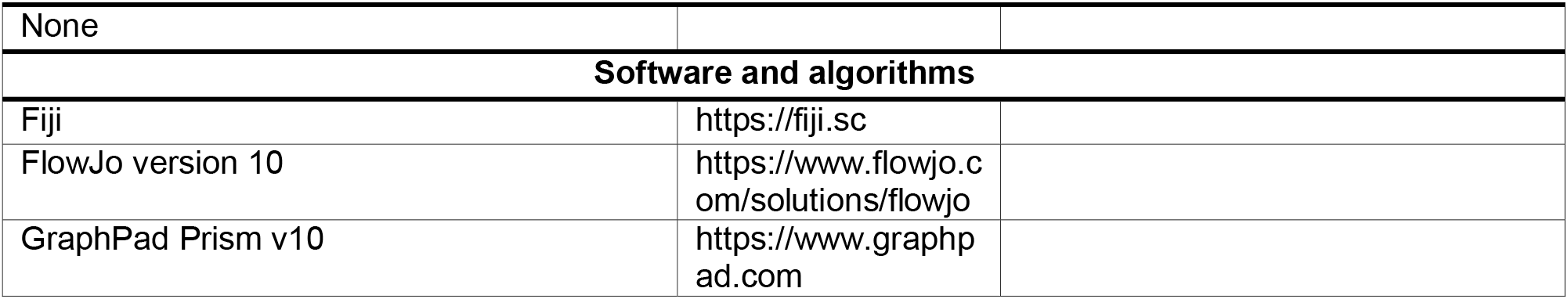

## Resource availability

### Lead contact

*Further information and requests for resources and reagents should be directed to and will be fulfilled by the lead contact, Shukti Chakravarti*.

### Materials availability

*All unique/stable reagents generated in this study are available from the lead contact*.

### Data and code availability

- All data reported in this paper will be shared by the lead contact upon request.
- This paper does not report any original code.
- Any additional information required to reanalyze the data reported in this work paper is available from the lead contact upon request.
- All raw bulk RNA-seq datasets will be deposited and made publicly available through the GEO database.

## Experimental model and study participant details

### Mice

All protocols were approved by the New York University Institutional Animal Care and Use Committee (IACUC). The C57BL/6J wild types (Jackson Laboratories, Stock#000664), *Bgn^KO^* (*Bgn^-/-^* and *Bgn^-/0^*)^58^ and *Lum^-/-^*^59^ all in the C57BL/6J background, were maintained in a specific pathogen-free mouse facility at NYU Grossman School of Medicine (NYU GSoM) in clear, air-filtered cages with 12-hour light/dark cycle and ad lib feeding. Mice, between the ages of 8-16 weeks, were used in all experiments, and the animals were sex-balanced within each experiment. All keratitis scoring was blinded. No animals were excluded unless they were sick or had ocular pathology prior to HSV-1 infection.

## Method details

### Cells, Viruses and Plaque Assay

HSV-1 McKrae and mCherry-expressing HSV-1 KOS strains were a kind gift from Drs. Daniel Carr (University of Oklahoma Health Sciences Center)^35^ and Prashant Desai (Johns Hopkins University School of Medicine)^38^. Viral strains were propagated in our lab using the Vero cells (Green African monkey kidney epithelial cells; ATCC#CCL81.2). Vero cells were grown in Eagle’s minimum essential medium (EMEM) supplemented with 10% fetal bovine serum (FBS) and antibiotic/antimycotic (Gibco). Virus stocks were prepared by infecting Vero cells at a multiplicity of infection (M.O.I.) of 0.01 PFU/cell. After 3 days post-infection, both intracellular and extracellular viruses were harvested, concentrated, and the purified virus was aliquoted and stored at -80 °C. The viral stock was titrated by standard plaque assay. Briefly, Vero cell monolayers were used to perform a plaque assay. The viral stock or HSV-1-infected corneal swabs were serially diluted 10-fold in Opti-MEM, and 100 µl of the viral dilution was overlaid on Vero cells in a 12-well plate for 1 h, with gentle rocking every 10 min. One-hour post-infection, Opti-MEM was replaced with a complete EMEM containing 2 % methylcellulose (Sigma-Aldrich). The plates were incubated at 37°C with 5% CO_2_ until visible plaques appeared (2-3 days). Methylcellulose was pipetted off, and cells were stained with 1% crystal violet (Sigma-Aldrich) in 50% ethanol for 20 min and washed with water to visualize plaque formation. The plaques were manually counted, and images were captured. The human telomerase reverse transcriptase-immortalized corneal epithelial cell line (hTCEpi), donated by James Jester, PhD (University of California, Irvine), was used between passages 78 and 82 ^60^. The hTCEpi cells were cultured in KGM Gold growth medium (Lonza) supplemented with antibiotics and antimycotics.

### UV-inactivation of HSV-1 and HTCEpi infection

The HSV-1 (McKrae) stock was diluted to 10^4^ pfu/ml in keratinocyte growth medium (KGM-Gold) medium (Lonza) and exposed to 395-nm UV light for 30 min in a UV chamber (Benchmark UV Clave). A 30 µl aliquot was taken after UV-inactivation for plaque assay to confirm viral inactivation. The telomerase-immortalized human corneal epithelial (hTCEpi) cells were grown in a 24-well plate containing KGM-Gold complete medium (500 ml of keratinocyte basal medium containing 30 µg/ml bovine pituitary extract, 0.12 ng/ml recombinant human epidermal growth factor, 5 µg/ml recombinant human insulin, 0.33 µg/ml hydrocortisone, 10 µg/ml recombinant human transferrin and 0.39 µg/ml epinephrine). hTCEpi cells were either pretreated with rLUM or rBGN for 60 min or left untreated, followed by infection with UV-inactivated HSV-1 (UV-HSV1) at a M.O.I of 1. For the positive control, hTCEpi cells were also treated with live HSV-1 at a M.O.I of 1. Cells were infected for 20 h at 37 °C in 5% CO_2_. The supernatant was harvested for CCL2 ELISA and the cell layer was washed and used for RNA extraction.

### Corneal infection, slit-lamp imaging and cornea scoring

8-16 weeks age, gender balanced C57BL6/J, *Bgn^KO^* and *Lum^-/-^* mice were anesthetized by intraperitoneal injection of 200 µl of ketamine and xylazine cocktail (10 mg/ml of ketamine and 0.66 mg/ml of xylazine in PBS). Complete anesthesia was confirmed by toe pinching. Once fully anesthetized, both corneas were scarified in a 3 x 3 grid using a 30-gauge needle. Following the corneal scratches, 4 µl of HSV-1 McKrae (10^4^ pfu) was added to the right cornea, while the scratched left cornea received PBS. The infected mice were housed in the ABSL2 pathogen-free animal facility. Mice were euthanized on 3 or 5 dpi and tissue was collected and processed for further analysis. Before euthanasia, mice were anesthetized and the eyes were scored in a blinded manner for corneal pathology and clouding by slit lamp biomicroscopy on 3 and 5 dpi. Slit-lamp images were used to score the corneal keratitis by two independent researchers in a blinded manner. The corneal opacity was scored on a four-point scale: 0.5, any corneal imperfection; 1, mild corneal haze; 2, moderate opacity; 2.5, moderate opacity with regional dense opacity; 3, diffuse dense opacity obscuring iris; 3.5, diffuse dense opacity with corneal ulcer; 4, corneal perforation^32^.

### Flow cytometry

The cornea and the submandibular lymph node were harvested from the mice after euthanasia. Corneas were treated with 3 mg/ml Collagenase type IV (Gibco) for 1 h at 37 °C in an orbital shaker (200 rpm). Similarly, the lymph nodes were treated with 25 µg/ml Liberase TM (Sigma) and 20 U/ml DNase I for 1 h at 37 °C. Single-cell suspensions were filtered through a 70 µm strainer, washed with ice-cold PBS, and stained with Live-Dead blue, followed by surface staining. Cells were stained in a 96-well V-bottom polypropylene plate. Cells were blocked with CD16/CD32 Fc block (1:200) in FACS buffer (PBS with 2% FBS and 5 mM EDTA) at 4 °C, then washed by centrifugation at 1500 rpm for 3 minutes at 4 °C. For surface staining, cells were stained with fluorophore-conjugated antibodies, washed 3x with FACS buffer, fixed in 4% paraformaldehyde (PFA), and resuspended in FACS buffer for analysis. All the gating was done on singlets and live cells. Fluorescence minus one (FMO) or isotype control was used as a negative control for gating. Multicolor flow cytometric analysis was performed on ZE5 (Yeti) analyzer (Bio-Rad) and analyzed using the FlowJo software (Tree Star, Inc.). All primary conjugated antibodies were used at a 1:200 dilution, and the antibody details are provided in the key resource table.

### ELISA and Multiplex bead-based cytokine analysis

Protein levels of human CCL2, LUM and BGN in the hTCEpi cell culture supernatant and infected organoid supernatant were measured using an ELISA kit according to the manufacturer’s instructions. Corneas from PBS- and infected-mouse groups were excised from the euthanized mice, and lysates were prepared using lysis buffer containing 0.1% TritonX-100, 50 mM Tris-HCl (pH 7.4), 150 mM NaCl, phosphatase inhibitors, and 1X protease inhibitor cocktail (Roche). Corneas were then homogenized, and the homogenate was centrifuged at 10,000 rpm for 15 min at 4°C. The clarified supernatant was collected, protein measured by Bradford assay and immediately frozen at -80°C for further analysis. 5 µg of protein/sample was used for the multiplex bead-based cytokine analysis according to the manufacturer’s instructions. The LEGENDplex mouse anti-virus response panel (BioLegend) was used to measure 13 pro-inflammatory cytokines and chemokines in corneal lysates, and the data were acquired on a BD FACSCanto flow cytometer. The LEGENDplex data analysis software was used to distinguish analytes based on bead size and internal dye.

### Bulk RNA-sequencing and analysis

Corneas were harvested from PBS and HSV-1-infected WT, Bgn^KO^ and Lum^-/-^ mice after euthanasia. The cornea was placed in an ice-cold tube containing RLT buffer, and the tissue was homogenized with a pestle. Total RNA was extracted using the QIAGEN RNeasy extraction kit (Qiagen) with DNase I treatment. RNA quantification, cDNA synthesis, fragmentation, library preparation (Illumina stranded mRNA v2 with polyA selection), and sequencing were performed by Genome Technology Center, NYUGSoM, using the NovaSeq X+ 10B platform. The raw read (FASTQ) files were checked for contamination, adaptor sequences, or other overrepresented sequences using FastQC 0.11.7; no contaminating sequences were found in the samples. Sequencing reads were aligned to the mouse reference genome (mm10) using STAR (v2.5.0c)^61^, with transcript annotations provided in GTF format. Insert size metrics were calculated using Picard tools (v1.126). Gene-level read counts were quantified using HTSeq (v0.6.0) (Anders et al., 2015). Raw count data were normalized on the basis of library size factors. Differential gene expression analysis between HSV-1 and PBS-treated samples was assessed separately within each genotype (WT, Bgn^KO^ and Lum^-/-^) using DESeq2 (v1.50.2) in R (v4.5.2). Genes with missing DESeq2 statistics were excluded prior to filtering. Differentially expressed genes (DEGs) were classified as significant according to the following criteria: FDR <0.05, log2 fold change > 1.5. To compare the HSV-1 versus PBS response across genotypes, differentially expressed genes (DEGs) lists were derived independently for WT, Bgn^KO^ and Lum^-/-^ and intersected to partition genes into the seven regions of a three-set diagram (genotype-exclusive, pairwise-shared, and common to all three). A proportional three-way Venn diagram was rendered with the VennDiagram R package. The ten most significant immune-related genes per plot were labeled with ggrepel; immune genes were defined as members of the Gene Ontology term "immune system process" (GO:0002376, including descendant terms; org. Mm. eg. db v3.22.0). Pathway enrichment analysis was performed using Gene Ontology Biological Process (GO BP) on the upregulated DEGs of each genotype using enrichGO in ClusterProfiler (v4.18.4) to analyze enriched pathways. Enrichment significance was tested with Fisher’s exact test. Sample relationships and replicate concordance were evaluated by principal component analysis (PCA) and Euclidean distance-based hierarchical clustering. All downstream statistical analyses and data visualization were performed in R (v3.1.1).

### Organoid infection

The human iPSC-derived corneal organoids (HCOs) were cultured and matured as previously described^36,62^. Around 3-month-old HCOs in a 96-well plate were infected with 1000 pfu of mCherry-expressing HSV-1 KOS strain for 1 h at 37 °C and 5% CO2, followed by three washes with 1x PBS, and then incubated for another 24 h. After infection, the supernatant was collected and stored at -80 °C for ELISA, and the HCOs were fixed with 4% PFA, followed by 20% and 30% sucrose, and OCT blocks were prepared and stored at -80 °C for histology.

## Quantification and statistical analysis

### Statistical Analysis

All the experiments were repeated two to four times, with results presented as mean ± SEM, as indicated in the figure legends. Comparisons between multiple groups were performed using one- or two-way ANOVA, followed by multiple comparisons analysis. Comparisons between the two groups were performed using a two-tailed unpaired Student’s t test with Mann-Whitney correction. Significance set at * p ≤ 0.05; ** p ≤ 0.01; *** p ≤ 0.001; ns, not significant. All statistical analyses were performed using GraphPad Prism version 10. All experiments were performed at least twice.

