## Supplementary data for "Extracellular matrix proteoglycans lumican and biglycan promote innate immune signals and viral clearance in HSV-1 infections of mouse corneas"

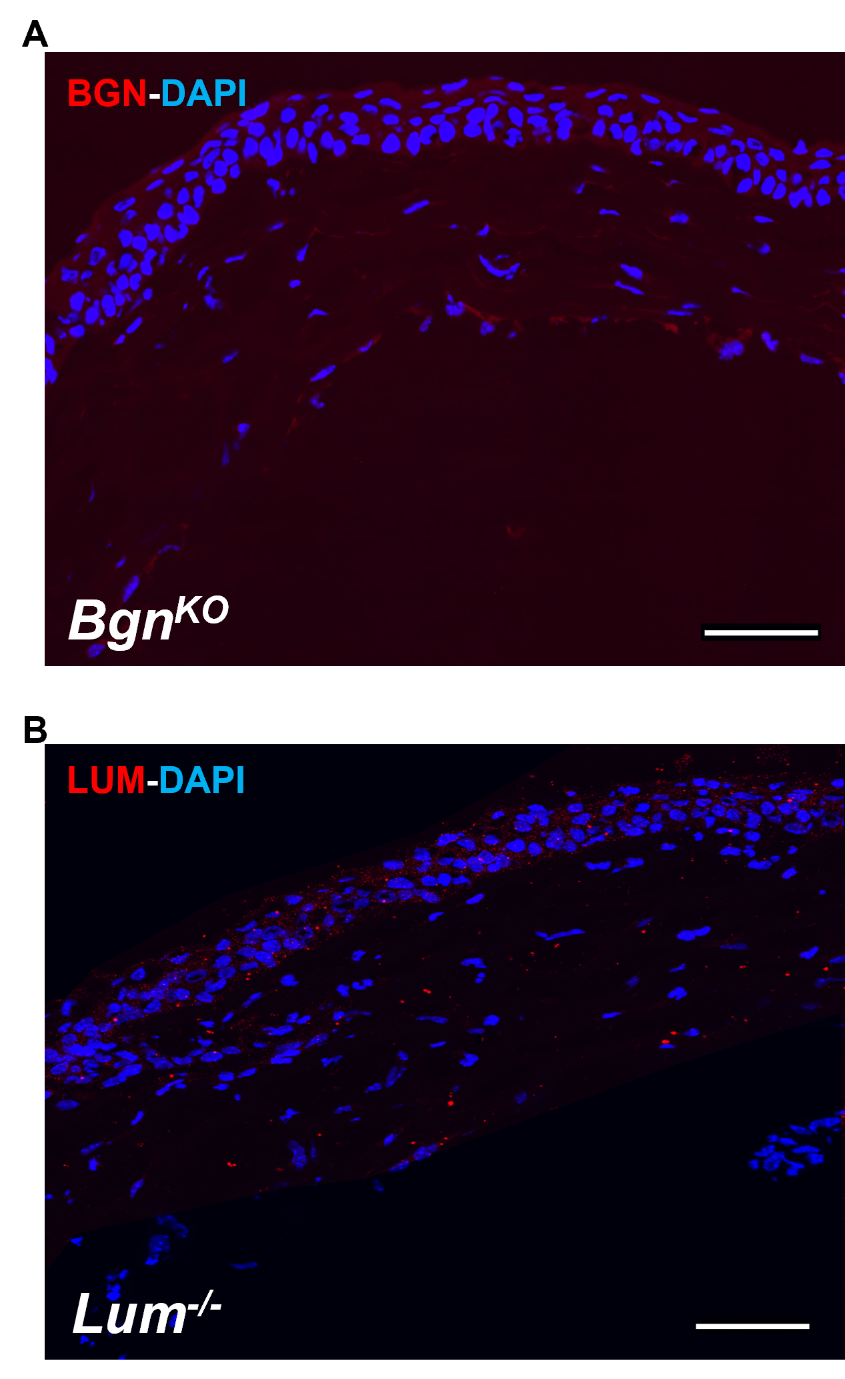
**Figure S1**

**Fig. S1. Immunohistochemistry of corneal cryosections** from *Lum^-/-^* (A) and *Bgn^KO^* (B) mice as controls for primary antibodies against Lum and Bgn showing minimal to no immunoreactivity. Scale bars: 100 µm.

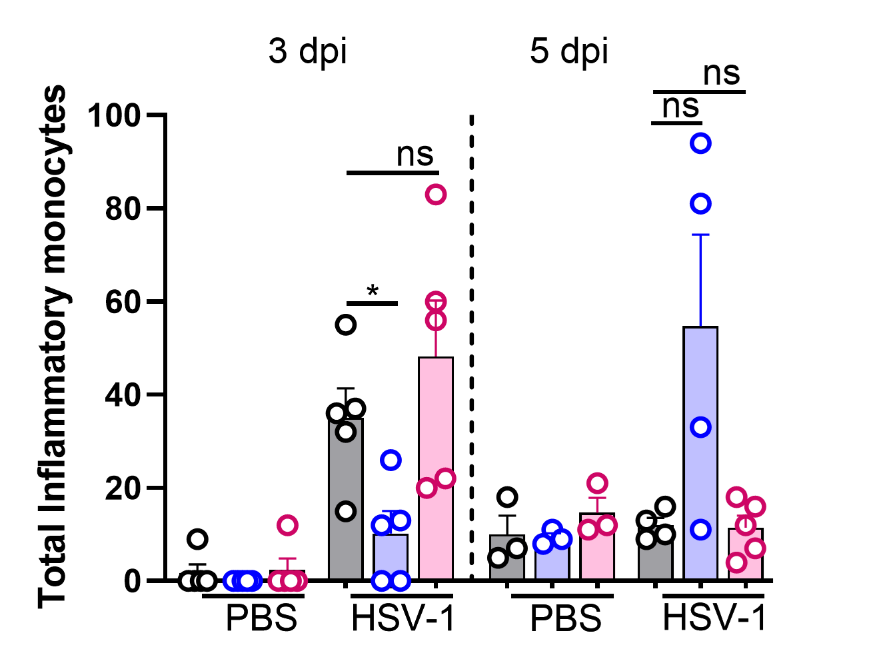
**Figure S2**

**Fig. S2. Inflammatory monocyte numbers in the HSV-1-infected mouse cornea.** Quantification of total inflammatory monocytes (CD11b^+^Ly6C^+^F4/80^+^) in corneas of data shown in Fig. 2C. Overall total inflammatory monocytes counts were very low. *Lum^-/-^* corneas showed significantly lower than that of WT at 3 dpi. By 5 dpi the total counts were similar in all three genotypes. n = 3 – 9 mice/genotype; *p < 0.05; ns, not significant. Statistical significance was calculated using an unpaired t-test with Mann-Whitney correction. Error bars represent the mean ± SEM.

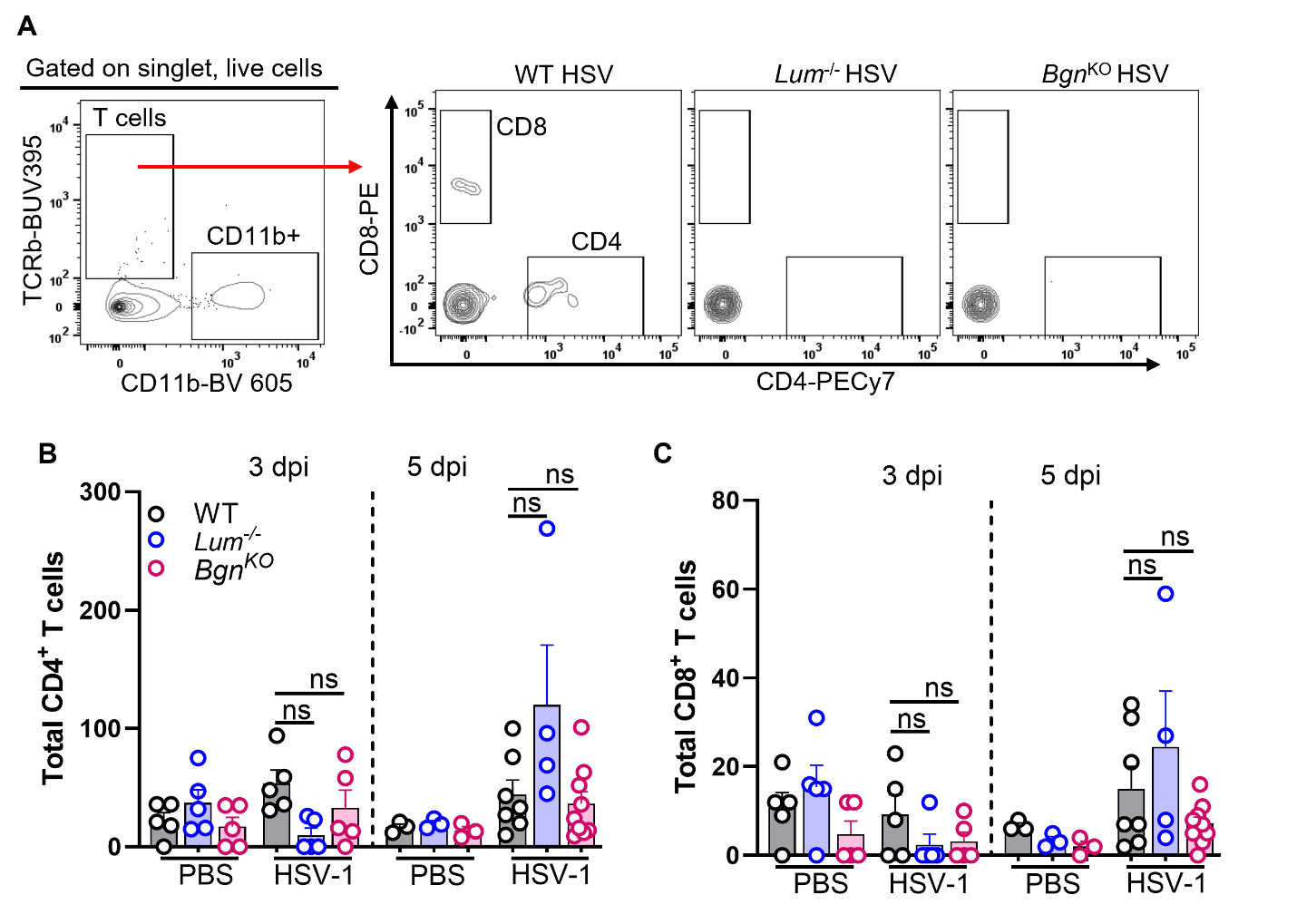
**Figure S3**

**Fig. S3. Lymphocyte numbers in the HSV-1 infected cornea are low and similar across all three genotypes.** Total CD4 (TCRβ^+^CD4^+^) and CD8 (TCRβ^+^CD8^+^) T cells in the corneas were quantified. n = 3 – 9 mice/genotype; ns, not significant. Statistical significance was calculated using an unpaired t-test with Mann-Whitney correction. Error bars represent the mean ± SEM.

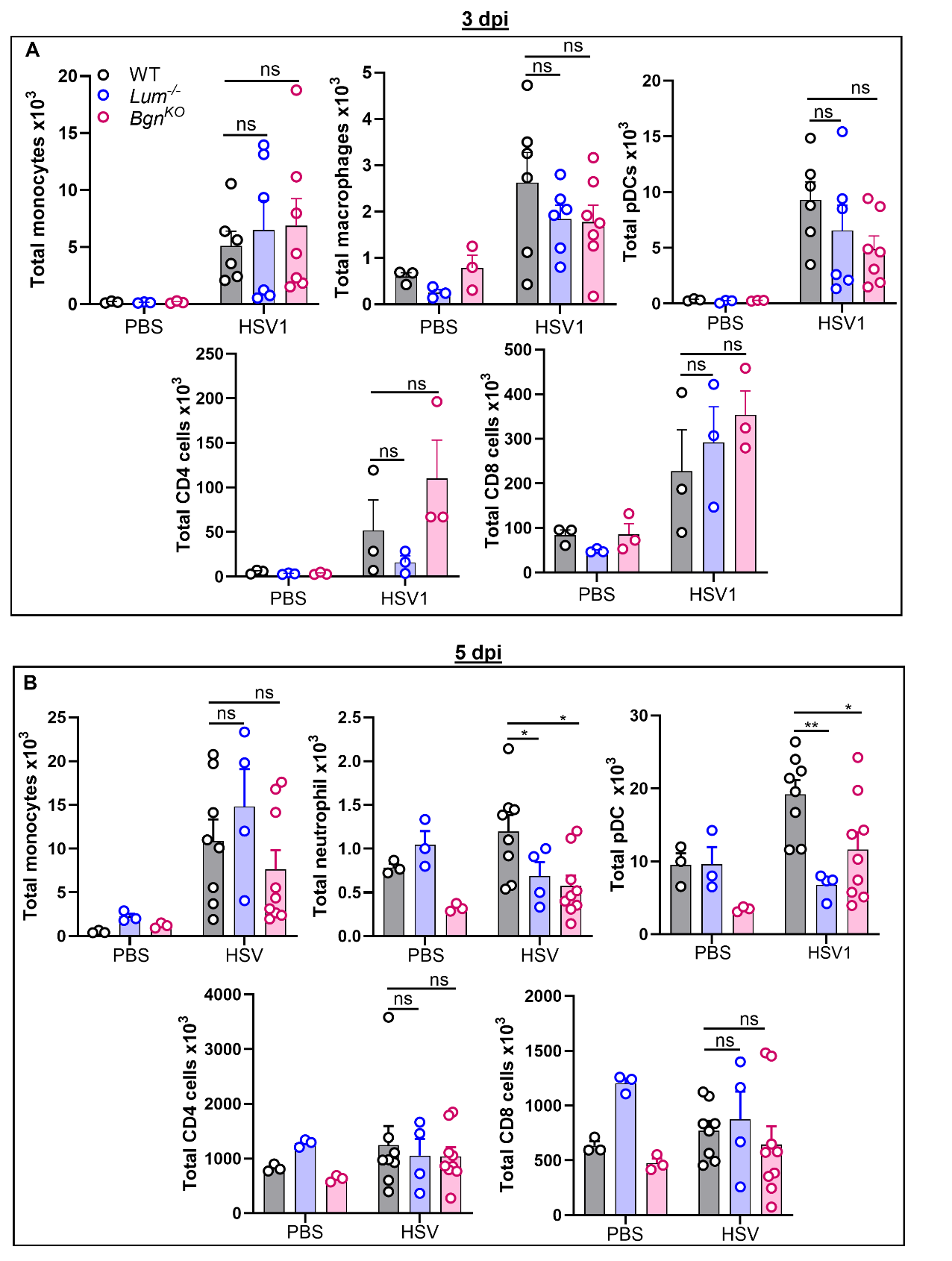
**Figure S4**

**Fig. S4. Myeloid and lymphoid cell numbers in the draining lymph node (dLN) of infected mice.** Myeloid cells were gated on singlets, live cells, and CD11b^+^ cells to enumerate monocytes (Ly6C⁺F4/80^-^Ly6G⁻), macrophages (Ly6C^-^F4/80^+^Ly6G⁻), neutrophils (Ly6C^-^Ly6G^+^) and pDCs (CD11c^+^B220^+^PDCA1^+^) in the dLN of HSV-1 infected and PBS-treated cornea. Lymphocytes were gated on singlets, live cells, and TCRβ^+^ cells to quantify CD4 (TCRβ^+^CD4^+^) and CD8 (TCRβ^+^CD8^+^) T cells. **(A)** At 3 dpi, the total number of myeloid and lymphoid cells was elevated in the dLN of infected mice compared to the PBS-treated control mice, but their numbers were similar in all three genotypes. (**B**) At 5 dpi, total number of neutrophils and pDCs in the dLN of infected mice were significantly higher in the WT compared to *Lum^-/-^* and *Bgn^KO^* mice. Lymphocytes number were comparable among the three genotypes. n = 3 – 9 mice/genotype; *p < 0.05; **p < 0.01; ns, not significant. Statistical significance was calculated using an unpaired t-test with Mann-Whitney correction. Error bars represent the mean ± SEM. Details of the corneal infections are provided in Fig. 2.

**Figure S5**

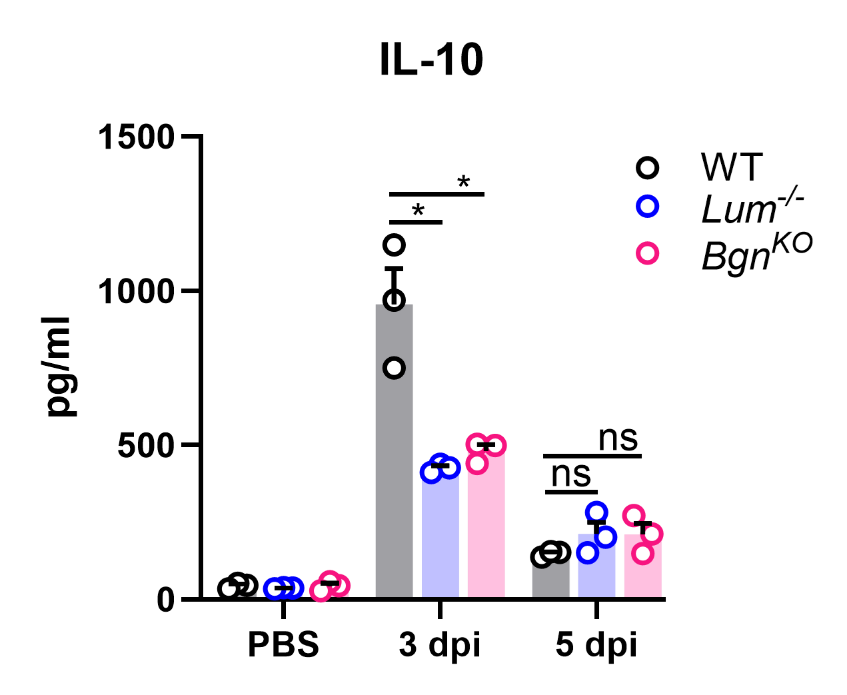

**Fig. S5. Anti-inflammatory IL-10 levels are higher in the infected WT corneas.** Anti-inflammatory IL-10 levels in the corneal homogenates were significantly elevated in HSV-1-infected WT corneas at 3 dpi compared to *Lum*^-/-^ and *Bgn^KO^* mice. By 5 dpi, IL-10 levels declined in WT mice and were similar to those in *Lum*^-/-^ and *Bgn^KO^* mice. n = 3 mice/genotype; *p < 0.05, ns, not significant. Statistical significance was calculated using unpaired t-test with Mann-Whitney correction. Error bars represent the mean ± SEM.

**Figure S6**

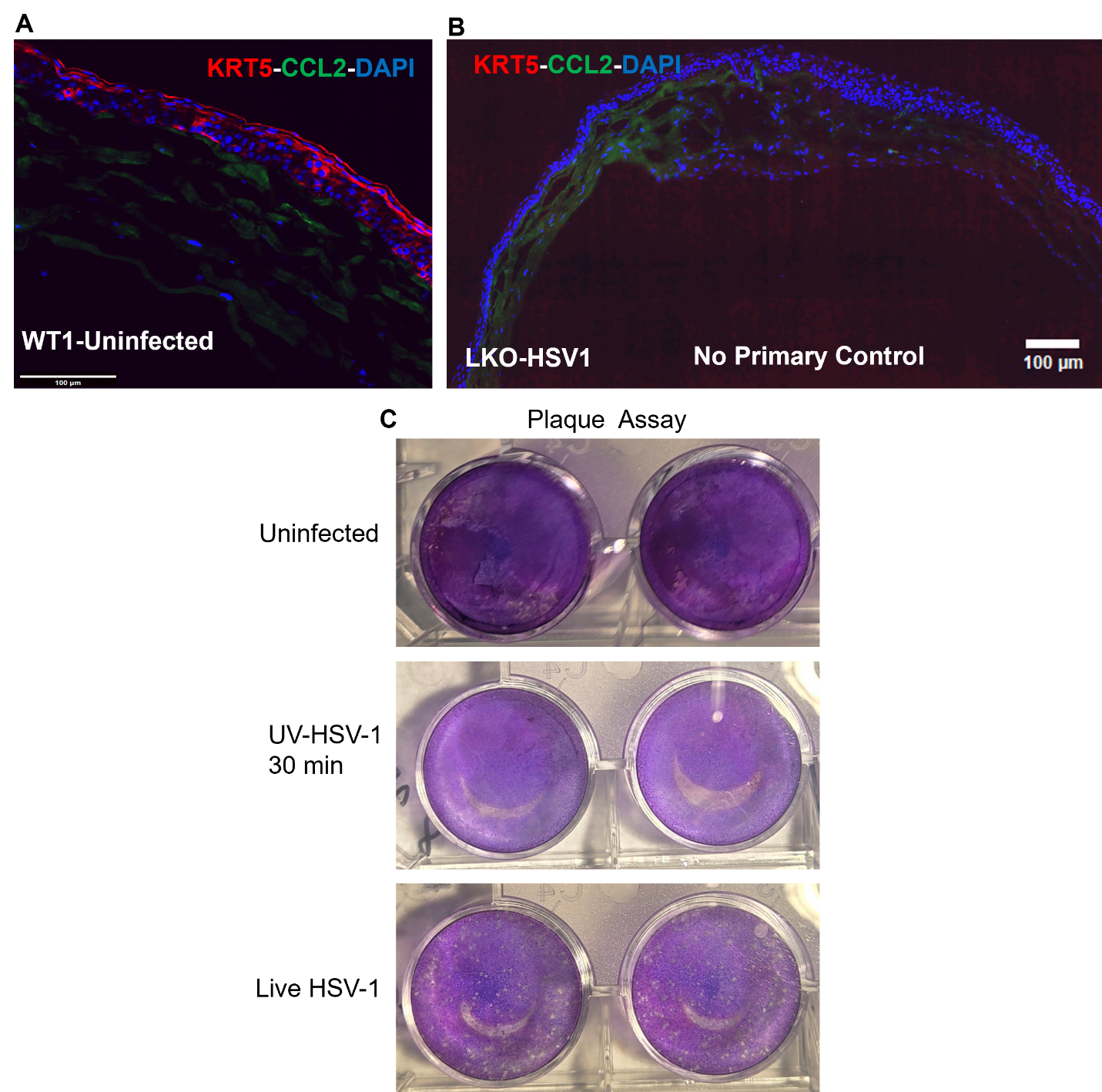

**Fig. S6. Uninfected WT corneas do not secrete Ccl2. (A)** Representative immunohistochemical staining of corneal sections from uninfected WT mice at 3 dpi demonstrates the absence of Ccl2 expression. Scale bars: 100 µm. One representative image from 2 mice. (**B**) Cornea cryosections from *Lum^-/-^* mice served as negative controls for the primary antibodies and showed minimal immunoreactivity. Scale bars: 100 µm. (**C**) Plaque assay images validate the inactivation of HSV-1 McKrae by UV exposure for 30 minutes.
